# Hippocampal epigenetic changes associated with population cycle phase in wild voles

**DOI:** 10.64898/2026.05.04.722675

**Authors:** Phoebe D. Edwards, Viswanathan Satheesh, Charles J. Krebs, Alice J. Kenney, Rudy Boonstra

## Abstract

Vole and lemming population cycles are an enigma in ecology. Decades of field observations and experimental manipulations have revealed that cycles cannot always be explained by extrinsic factors in the environment, including food availability or predator numbers. Thus, it has been proposed that intrinsic mechanisms, such as adaptive alterations in phenotype during different phases of the cycle, drive population dynamics. However, the mechanisms underlying such phenotypic changes have not been elucidated. We test the hypothesis that epigenetic changes occur over population cycles by comparing whole epigenome DNA methylation changes in brain tissue collected from northern red-backed voles (*Clethrionomys rutilus*) in a wild, naturally cycling population during the peak, decline, and low years. Overall, the greatest number of differentially methylated CG sites (DMCs) and differentially methylated regions (DMRs) were detected in comparisons between voles from the peak phase and low phase of the cycle. We highlight methylation differences in the promoter region of ATP synthase subunit c (*Atp5g3*) and an intron of insulin-like growth factor 1 receptor (*Igf1R*), which may be associated with growth, development, and bioenergetics. There were additional changes in the promoters of members of the cytochrome P450 enzyme family, including *Cyp1a1*, associated with estrogen metabolism, as well as the promoter of macrophage migration inhibitory factor (*Mif*), and in an exon of serum/glucocorticoid regulated kinase (*Sgk1*), which may link changes in stressors to direct brain changes. Our study is the first interrogation into broad epigenetic changes associated with natural population cycle phase in a wild mammal.

## 1 Introduction

Population cycles in voles and lemmings have been the subject of intensive ecological research for over 100 years. Across many species, and in many regions of the world, these rodents exhibit pronounced changes in population density over 3-5 year periods (reviewed in Krebs 2013). Cycles are characterized by repetitive periods of population increase, peak, decline, and low phases, with each phase lasting approximately one year. Because voles and lemmings are a food resource for many predators, changes in their numbers have ramifications for other aspects of the ecosystem, including the presence, numbers, and reproductive output of terrestrial and avian predators (Gilg et al. 2006; Côté et al. 2007; Giraudoux et al. 2020). These cycles can also influence the available plant biomass, with high densities sometimes reducing plant biomass (Olofsson et al. 2012). Much work has addressed the factors that may drive these population cycles, especially the influence of predator numbers and food availability, and has revealed many examples where changes in these factors are not able to explain population cycles (Cole et al. 1978; Maron et al. 2010; Bolduc et al. 2025). In one particular example, a large-scale experiment in the boreal forest of Yukon, Canada, manipulated terrestrial predator presence, food availability, nutrient availability (fertilizer), and various combinations of these treatments, across large plots for nine years (1988-1996). However, none of these treatments were able to significantly change northern red-backed vole (*Clethrionomys rutilus*) population density relative to control sites (Boonstra & Krebs 2006). We reassessed this data in 2025 using capture-mark-recapture models and found that the original conclusions were upheld; any increase in vole recruitment was offset by poorer apparent survival, and vice versa (Edwards et al. 2025).

Because of the insufficient explanatory power of factors like food and predation in many of these cases, intrinsic regulation (or population self-regulation) has been proposed as a driver for these cycles (Chitty 1960; Boonstra et al. 1998). Intrinsic regulation refers to population regulation by factors internal to the population, such as changes in compositions of genotypes and phenotypes, animal quality (e.g. maternal quality, offspring quality), behavior, or dispersal patterns (Mihok & Boonstra 1992; Ostfeld et al. 1993). The earliest iteration of this hypothesis was Chitty’s 1967 proposal that at high population densities, natural selection favored a high-aggression, low-reproduction genotype, initiating the decline. Once population densities were low, selection for a high-reproduction, low-aggression genotype would reoccur (Chitty 1967).

Indeed, there is evidence that voles at different phases of the cycle are phenotypically different: in the peak phase of the cycle, when densities are highest, a smaller proportion of the adult population is reproductive, there is delayed sexual maturation of juveniles (Krebs et al. 1969; Boonstra 1985; Norrdahl & Korpimaki 2002, Novikov et al. 2012) and sometimes very large individuals are observed (~25% larger than the average body mass), termed “the Chitty effect” (Chitty & Chitty 1962; Oli 1999; Olea et al. 2024). Chitty’s hypothesis of rapid selection to produce these alternate morphs at high and low density has been rejected as captive-breeding studies of voles and lemmings have demonstrated low paternal heritability of reproductivity, growth rate, and body mass (Boonstra & Boag 1987; Boonstra & Hochachka 1997). An alternate mechanism that may facilitate cycle-related changes in animal quality is early-life epigenetic change (e.g. changes in DNA methylation) in response to population density conditions (Boonstra 2024). In this way, early-life or maternal conditions could modify long-term phenotype, potentially in an adaptive way in response to phase-specific social conditions.

To test for population cycle-related differences in DNA methylation in the brain, we collected tissue from Northern red-backed voles at a long-term study site in the boreal forest in Yukon, Canada, where they have been monitored for 50 years. Red-backed voles, like many other voles, have a social system of promiscuous breeding and female territoriality (Ostfeld 1985; Krebs et al. 2007). During the reproductive season, which lasts from approximately May to September, adult female voles establish exclusive territories, wherein female-female spatial overlap is low (Bujalska 1973; Gilbert et al. 1986; Boonstra & Krebs 2012). The offspring born that spring and summer (the “young of the year”) can either become reproductive that breeding season or delay reproduction until the following year. The latter strategy of delayed reproduction is more common when population densities of reproductive adults are already high (Boonstra 1978; Saitoh 1981; Gilbert et al. 1986; Edwards et al. 2021). In northern red-backed voles, overwinter survival is poor for the breeding adults, hence the maximum lifespan is around one year (Boonstra & Krebs 2012). Thus, there is nearly complete turnover of the breeding population each year.

We assessed whole-epigenome methylation changes in hippocampal brain tissue collected from female young of the year voles during fall live-trapping in peak, decline, and low phases of the population cycle. From a practical standpoint, we focused our analysis on a single sex and age/reproductive class to reduce epigenetic variation that may naturally occur between sexes and ages/reproductive statuses. In terms of population cycle-related changes in physiology, we expect that females are key limiting factors for population growth, due to female spacing behavior and because one male vole can impregnate many females, but females are constrained by the investments of pregnancy and maternal care. Further, prior work in other vole populations has shown that females can have more pronounced changes in gene expression in the brain in response to population density conditions (Edwards et al. 2021; Edwards et al. 2023). We focused on the hippocampus in the brain, because this region has been well-studied in terms of epigenetic changes in response to early-life conditions and stressors in rodent models (Weaver et al. 2004; McEwen 2010).

## 2 Materials and Methods

### 2.1 Population monitoring and tissue collection

We studied red-backed vole populations in the North American boreal forest near Kluane Lake in the southwestern Yukon (60°57′ N, 138°12′ W). Small mammals at this site have been monitored since 1973 (Studd et al. 2025). Small mammals are live trapped twice a year using Longworth live traps, in late May and in late August-early October. Population density is determined using the average of three 10 x 10 control grids, with a 15.24 m interval between traps. Density estimates are made with Efford’s maximum likelihood estimator. Further details on live-trapping procedures at this long-term monitoring site can be found in prior publications (Boonstra & Krebs 2006, Krebs et al. 2023).

We collected voles during fall trapping in 2017, 2018, and 2019, representing population peak, decline, and low phases, respectively (**Figure 1**). A subset of live trapped animals was brought to the laboratory at Kluane Lake Research Station (approximately a 10 min drive from the field site). Voles selected were non-reproductive, juvenile females, as determined by lack of vaginal perforation, lactational tissue, and of implantation sites in their uteri. Based on the population turnover rate in this species and the range of body masses in collected animals (11-23 g), we estimated that these animals were young of the year. Voles were overdosed with isoflurane and rapidly decapitated. Brain tissue was removed and immediately frozen on dry ice. Tissue was subsequently stored at -80° C until microdissection at the University of Toronto Scarborough.

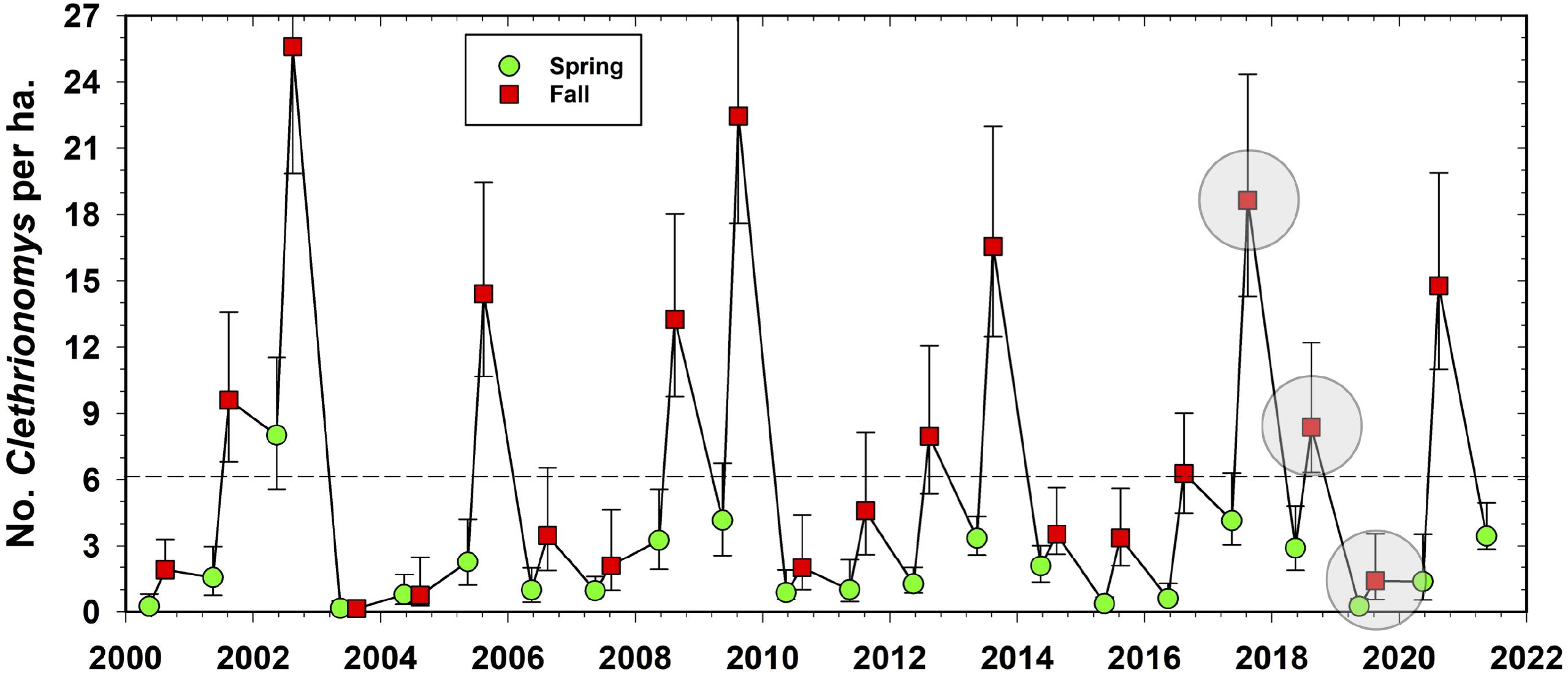

Brains were dissected into 30 um microsections using a cryostat (Leica CM3050S, Leica Biosystems), targeting the hippocampus based on brain landmarks in the Allen Mouse Brain Atlas (Lein et al. 2007). DNA was then extracted from hippocampal tissue with the MasterPure Complete Purification Kit (Epicentre). All animal work was done in accordance with the guidelines of the Canadian Council on Animal Care and approved by the Animal Care Committee at the University of Toronto.

### 2.2 Enzymatic methyl-sequencing and data analysis

All library preparation and sequencing was conducted at the Centre for Applied Genomics (TCAG), Hospital for Sick Children, Toronto. Hippocampal DNA from a total of 25 individual voles (8 from the peak phase in 2017, 8 from the decline phase in 2018, and 9 from the low phase in 2019) was sent for library preparation using the NEB Next Enzymatic Methyl-Seq kit (New England Biolabs). Libraries were sequenced using Illumina NovaSeq, with 150 bp paired end reads, resulting in an average of 171.5 million reads per sample (see Supporting Information, Table S1).

Because both EM-seq and bisulfite sequencing result in conversion of unmethylated cytosines, tools and pipelines developed for bisulfite sequencing can be used for EM-seq data (Olova & Andrews 2025). Reads were trimmed using Trim Galore! (Krueger 2015) based on recommendations for EM-seq (Olova & Andrews 2025): --clip_R1 10 --clip_R2 10 – three_prime_clip_R1 10 --three_prime_clip_R2 10. Trimmed reads were aligned to the northern red-backed vole genome (GCA_040207285.2) that had been bisulfite converted using the package Bismark (Krueger & Andrews 2011). Alignments were deduplicated and methylation calling was performed with Bismark. Principal components analysis of cytosine methylation profiles across the genome were visualized with MethylKit (Akalin et al. 2012).

Genetic structure of the population can be a confounding factor in epigenetic studies; if related animals are sampled within a given condition, methylation data may show trends driven by relatedness rather than environmental factors (Lea et al. 2017; Laine et al. 2023). To determine relatedness in the voles in this study, we used BS-Snper (Gao et al. 2015) which is designed for extracting SNP data from bisulfite converted DNA. We then constructed a relatedness matrix from SNP data using SNPRelate (Zheng et al. 2012) and determined there was no relatedness structure among the individuals sampled (Supporting Information, Figure S1), i.e. voles collected for brain tissue were indeed randomly sampled from the general population and results should not be biased by relatedness. As no relatedness structure was detected in the population, analysis of differentially methylated CG sites and regions by cycle phase was conducted with the package DSS (Park & Wu 2016; Feng & Wu 2019) which uses a beta-binomial model that accounts for overdispersion. The threshold for differentially methylated CG sites was set at FDR < 0.05 and the criteria for DMRs was specified as regions with differentially methylated cytosines sites at a threshold of p < 0.01, following the p value threshold parameters specified in a recent study in mammalian brain tissue (Yuan et al. 2022). All other parameters for DMR inclusion were kept as the default parameters in DSS (min length = 50 bp, min CG = 3, dis.merge = 50 bp, pct.sig = 0.5). Models were run in R version 4.4.0 (R Core Team 2024).

DMCs and DMRs were linked with their associated genes and genomic elements using the package Genomation (Akalin et al. 2015). Promoter regions were considered to be regions ≤2 kb upstream of the transcription start sight. Intergenic regions were considered to be regions >2 kb from any gene.

## 3 Results

Principal components analysis indicated that voles did not cluster by cycle phase (Supporting Information, Figure S2), indicating there was no overall epigenetic signature of cycle phase. However, there were many specific changes in CG methylation by cycle phase, as demonstrated by the beta-binomial models. Between peak phase and decline phase voles there were 747 differentially methylated CG sites (DMCs) and 357 differentially methylated regions (DMRs). Between decline phase and low phase voles there were 853 DMCs and 439 DMRs.

Low phase and peak phase voles had the most differences, with 1,367 DMCs and 467 DMRs. When examining the genomic context of the DMCs, an average of 0.9% DMCs across the three comparisons occurred in the promoter region of genes, 33.3% occurred in introns, 1.5% in exons, and 63.2% in intergenic regions. Similarly, of the DMRs, an average of 1.0% occurred in promoters, 37.1% in introns, 2.9% in exons, and 57.8% in intergenic regions (Supporting Information, Figure S3). Of the genes associated with DMCs and DMRs, 95.8% and 95.0%, respectively, were annotated (gene identity at the location in the genome is known).

Focusing on methylation differences in promoter regions, there were 27 DMCs and 13 DMRs overall (Table 1). Cycle phase comparisons showed DMCs in the promoter of the gene for a subunit of ATP synthase (*Atp5g3*), including two sites (909 and 910 bp from the TSS) hypomethylated in the peak relative to the decline, and one site (910 bp from the TSS) hypomethylated in the low relative to the decline (**Figure 2A**). There were additionally multiple instances with DMRs in the promoter region of cytochrome P450 enzymes; there was a 52 bp region in the promoter of cytochrome P450 1A1 (*Cyp1a1*) hypermethylated in decline relative to low phase voles, and a 305 bp region in the promoter of cytochrome P450 family 2 (*Cyp2j13*) hypermethylated in peak relative to decline phase voles (**Figure 2B**). There was a 73 bp DMR in the promoter of macrophage migration inhibitory factor (*Mif*) that was hypomethylated in low phase voles relative to decline phase (**Figure 2C**).

**Table 1.**
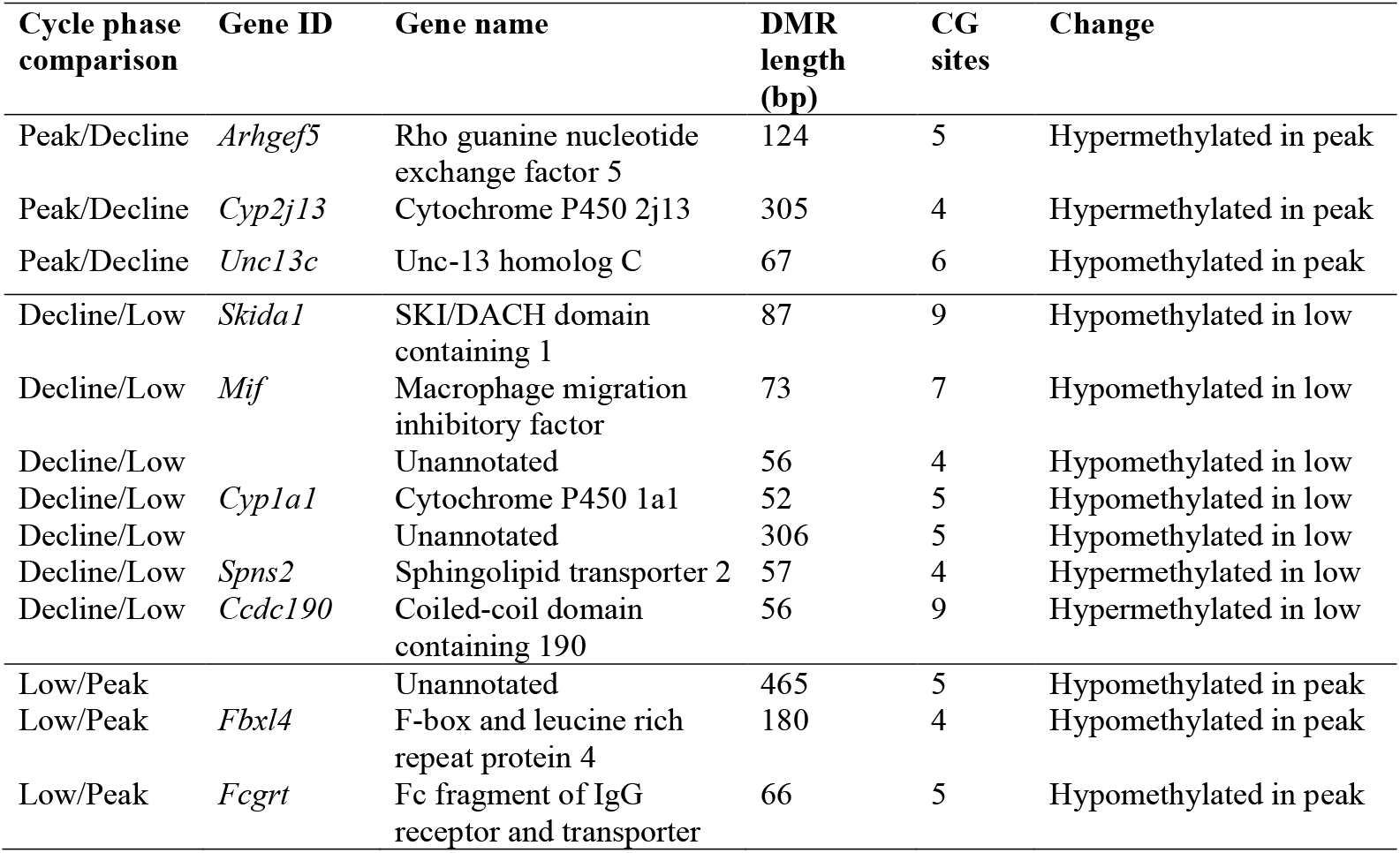
Differentially methylated regions (DMRs) in gene promoters.

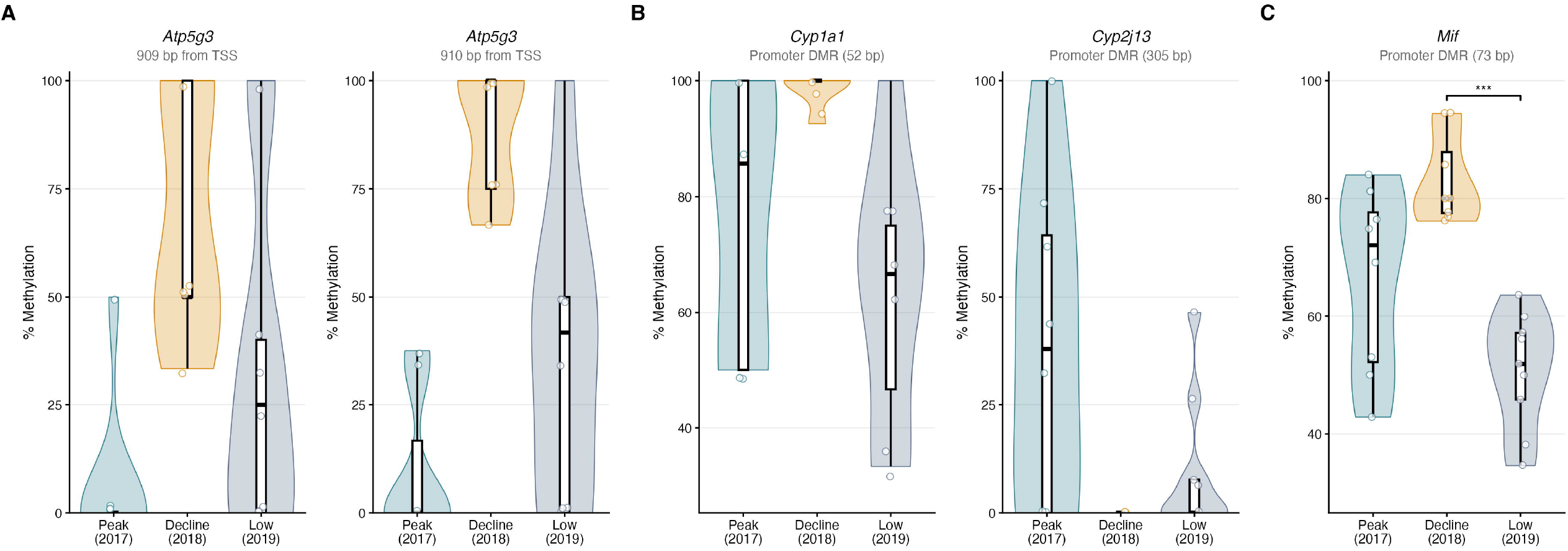

Within gene bodies, there was a 53 bp DMR in an exon of serum/glucocorticoid regulated kinase 1 (*Sgk1*), as well as a 66 bp DMR between peak and low phase voles, but no methylation differences in *Sgk1* between peak and decline phase voles (**Figure 3A**). There was a 72 bp DMR in the insulin-like growth factor 1 receptor (*Igf1R*) intron between peak and decline phase voles, a 95 bp DMR in the *Igf1R* intron between decline and low phase voles and between peak and low phase voles (**Figure 3B**), as well as a 59 bp DMR in the intergenic region closest to IGF-1 binding protein (*Igf1bp7*).

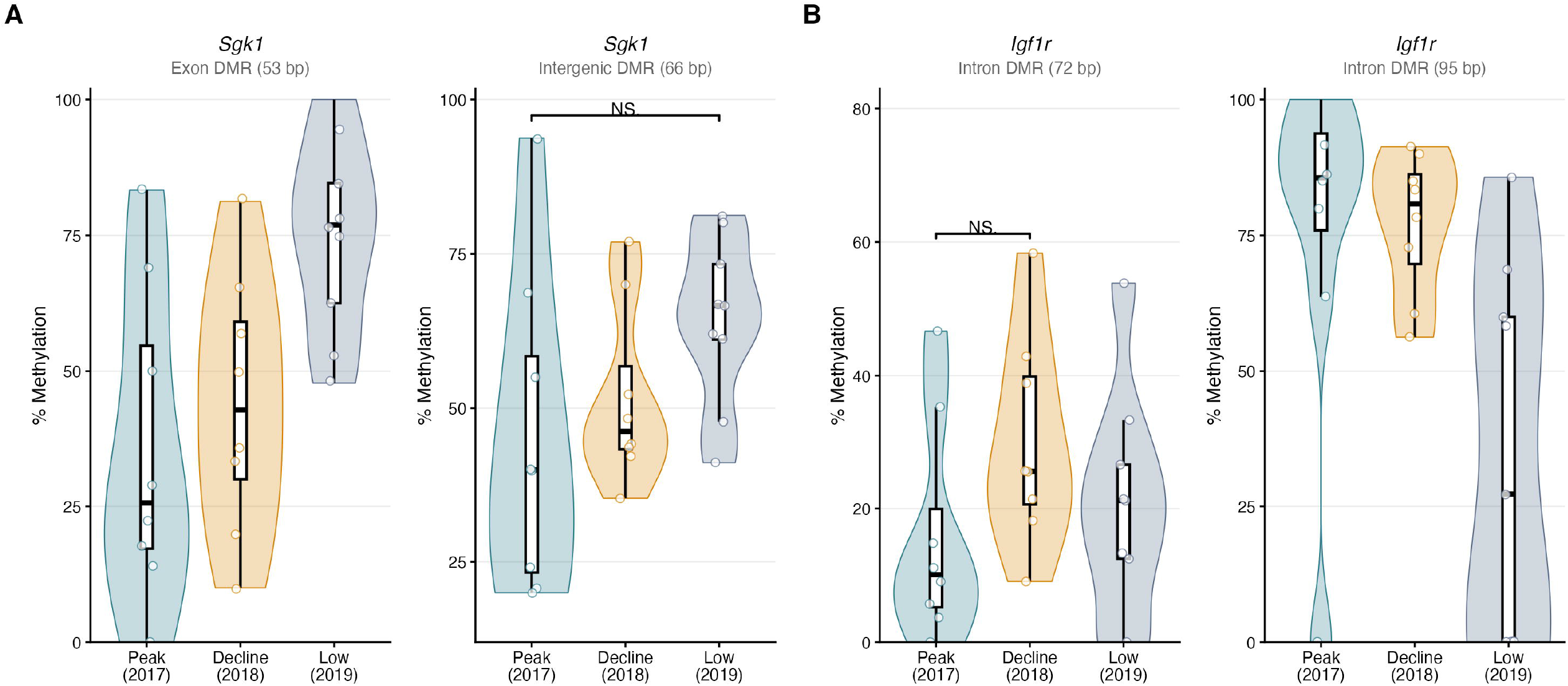

The majority of DMLs and DMRs were in intergenic regions of the genome. While it is unclear what the implications of these differences in methylation imply for gene expression, we highlight a few additional intergenic regions where the most closely located gene is of functional interest. There were multiple DMRs most closely associated with estrogen-related receptor gamma (*Esrrg*), including three DMRs between decline and low phase voles, one between peak and decline phase voles, and one between peak and low phase voles. Neuropeptide Y (*Npy*) was the gene most closely associated with three DMRs (237 bp, 97 bp, 65 bp) between peak and low phase voles; similarly, was an additional 162 bp DMR closest to neuropeptide receptor Y5 (*Npy5r*) between peak and low phase voles. Between decline and low phase voles, there was a large, 466 bp DMR closest to the *Npy5r* gene. There were two DMRs (59 bp and 80 bp) in the intergenic region closest to *Atp5g3* between decline and low phase voles, and a 109 bp DMR in the intergenic region between peak and decline phase voles. The full list of DMCs and DMRs in all regions can be found in the Supporting Information.

## 4 Discussion

Red-backed vole juveniles collected from different phases of the population cycle had numerous DMCs and DMRs, with the greatest number of overall differences occurring between the peak and low phase voles. Yet, within promoter regions specifically, most differentially methylated regions were observed in the decline voles relative to the other phases (**Table 1, Figure 2**). Although this study does not directly measure gene expression, we can make predictions about expression differences associated with some of these differentially methylated sites and regions. The most common DNA modification is the 5-methylcytosine (5mC) mark, and these methyl marks have generally (but not always) been associated with decreased gene expression, particularly when they occur in promoter regions, where they can interfere with transcription factor binding (Suzuki & Bird 2008, Kang et al. 2019). The effects of methylation in gene bodies have been associated with both decreased and increased gene transcription, potentially in a target specific/region specific way (Seigfried & Simon 2010; Hahn et al. 2018).

One of the key methylation differences that emerged in gene promoters was increased methylation of sites in *Atp5g3* in decline phase voles (**Figure 2A**), which we hypothesize could result in lower expression of ATP synthase subunit c during the decline. If ATP synthase production is affected by these modifications, this would suggest that cellular metabolism is reduced in decline phase voles (Vives-Bauza et al. 2010; Spataru et al. 2019). Decline phase voles additionally had DMRs in the promoters of two genes encoding cytochrome P450 enzymes. A DMR in the *Cyp1a1* promoter indicated hypermethylation in decline voles (**Figure 2B**), suggesting decreased production of the enzyme CYP1A1. This enzyme metabolizes estradiol; mRNA expression of *Cyp1a1* is markedly higher in the tissues of female laboratory mice compared with males, and this difference has been directly linked to promoter methylation differences between the sexes (Penaloza et al. 2014). Potentially lower *Cyp1a1* expression in the female decline phase voles may therefore indicate less active estradiol production, aligning with observations that reproduction is supressed during the decline phase. A DMR in the *Cyp2j13* promoter was hypomethylated in decline voles (**Figure 2B**), suggestive of increased expression of *Cyp2j13*. In laboratory mice, the CYP2J family has been linked to fatty acid and lipid metabolism, as well as the breakdown of xenobiotics (Graves et al. 2013), but there is generally little information about this family and the functional role of the J13 isoform. Changes to insulin-like growth factor 1 (IGF-1) signaling are of interest with respect to physiological changes underlying population cycle phase. IGF-1 is a hormone largely produced by the liver but can cross the blood-brain barrier and is additionally produced locally in the brain (Nieto-Estevez et al. 2016). Throughout tissues in the body, IGF-1 is associated with growth; knockouts or mutations of IGF-1 and IGF-1 receptor (IGF-1R) result in impaired growth, while in the brain, IGF-1 signaling has been implicated in hippocampal neurogenesis and other neuroprotective functions such as inhibition of apoptosis (Nieto-Estevez et al. 2016). A DMR within an intron of *Igf1R* was hypomethylated in voles in the low phase, albeit with high variation among individuals (**Figure 3B**). We hypothesize that IGF-1 signaling may be altered over cycle phases.

The potential role of chronic stress during population cycles has been an area of interest in rodents and other species. However, measurement of basal glucocorticoid (corticosterone) concentrations across vole population cycles or under different population density conditions have produced inconsistent results (reviewed in Edwards et al. 2023). Despite this, it is possible that population cycle conditions alter aspects of the stress response or stress-related pathways in the brain. Some results of this study can be related to this idea. Macrophage migration inhibitory factor (MIF) is a cytokine secreted by neurons that signals neuroinflammation and activates microglia (Zeng et al. 2024). Stimulation of the stress response and glucocorticoids are associated with increased MIF secretion (Calandra et al. 1995; Vedder et al. 2000). Serum/glucocorticoid regulated kinase 1 (SGK1) is an enzyme which catalyzes the phosphorylation of a variety of proteins and thereby plays a role in numerous physiological processes including cell survival and proliferation, ion transport, and immune function (Jang et al. 2022). The glucocorticoid receptor is one of multiple transcription factors that regulates SGK1 expression. Experiments in laboratory rodents and in human cell-cultures indicate that SGK1 increases in hippocampal tissue in response to glucocorticoid treatment and prenatal/early life stress (Millette et al. 2025). Interestingly, both elevated MIF and SGK1 are studied in models of human depression (Edwards et al. 2010; Millette 2025). Both of these targets were differentially methylated in low phase voles: a hypomethylated DMR in the promoter of *Mif* in low phase voles (**Figure 2C**) and a hypermethylated DMR in an exon of SGK1 (**Figure 3A**). If the functions of MIF and SGK1 in these wild rodents are aligned with the studies in human and laboratory rodent biomedical models listed above, this could be suggestive that animals born in the low phase may be experiencing stressors that are unrelated to spacing behavior and intraspecific competition (which would be most intense during the peak and decline, and relaxed during the low). Such stressors during the low could perhaps be related to delayed-density dependent changes in plant defenses/quality (Reynolds et al. 2012) or increased vulnerability to predation.

A caveat of our study is that it is a snapshot of methylation differences that occurred over one population cycle and does not replicate these differences across multiple cycles. The demographic pattern we observed (**Figure 1**) was a replicate of previous cycles over the last 50 years (Krebs et al. 2023; Studd et al. 2025), but there could be unknown environmental changes that occurred during one of the years sampled here that could influence the epigenome beyond the effects of cycle phase. However, these data are valuable as the first whole-epigenome data collected across a small mammal population cycle. While research in laboratory rodents is foundational in understanding the function of the genes listed above, we have little understanding of their patterns and functions in wild systems. Future work can directly compare commonalities or differences in other rodent cycles to better understand the generality of these changes. Further, epigenetic changes in other tissue types beyond the hippocampus are of interest. In a previous study on voles in experimentally manipulated population densities, we examined candidate gene methylation and expression in the hypothalamus of the brain and detected increased methylation of CG sites in the promoter region of the gene encoding gonadotropin releasing hormone (*Gnrh1*), a major component of the vertebrate reproductive axis (Edwards et al. 2021). In our current study, we did not interrogate tissues relevant to the reproductive axis but expect that this would be an area of change more broadly across population cycles (Lambin et al. 2025). Promoter methylation differences in *Cyp1a1* are promising that broader reproductive axis differences exist among population cycle phases. This work sets the stage for comparison with future studies and provides key insights into whole-epigenome data about underlying physiological changes that occur in voles during a natural population cycle.

## Author contributions

R.B. and P.D.E. conceptualized the project. B.S.G., C.J.K., A.J.K, and R.B. conducted field investigations. P.D.E. and V.S. conducted formal analysis and performed data visualization. P.D.E. and R.B. wrote the original draft. All authors reviewed and edited the manuscript.

## Acknowledgements

We thank the Champagne and Aishihik First Nations and Kluane First Nation for allowing us to work within their traditional territories. We thank Sophia Lavergne and Coral Frenette-Ling for assistance with brain tissue microdissection and DNA extraction.

## Funding

This work was supported by NSERC to R.B. and by Iowa State University support to P.D.E.

## Data Availability Statement

Associated data is available at https://doi.org/10.25380/iastate.32056107

The raw sequencing data can be found in the NCBI Sequence Read Archive under accession PRJNA1461410.

## Conflict of interest statement

The authors declare no conflicts of interest.

## Notes

### Competing Interest Statement

The authors have declared no competing interest.

