## Supplementary figures and images for "Hippocampal epigenetic changes associated with population cycle phase in wild voles"

### Figure S1

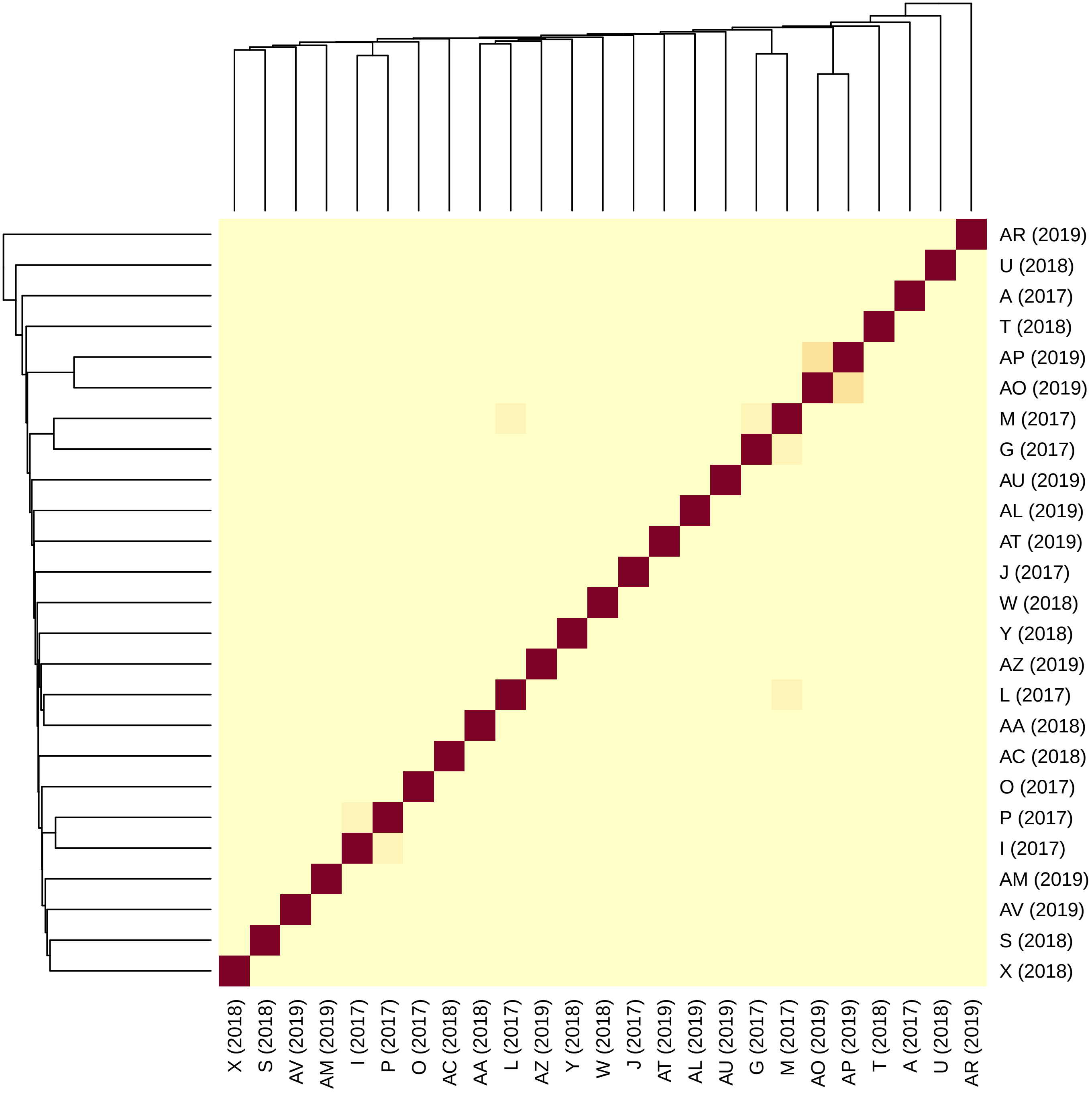

### Figure S2

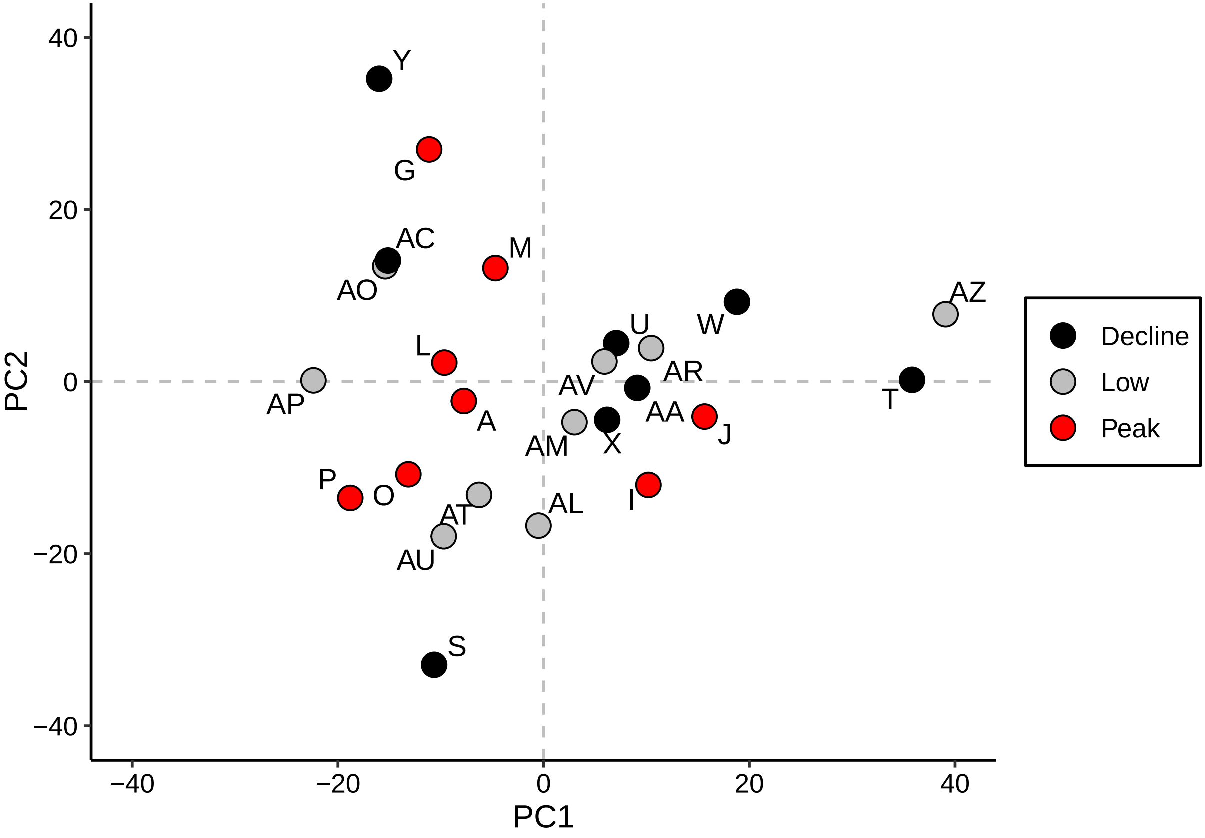

### Figure S3

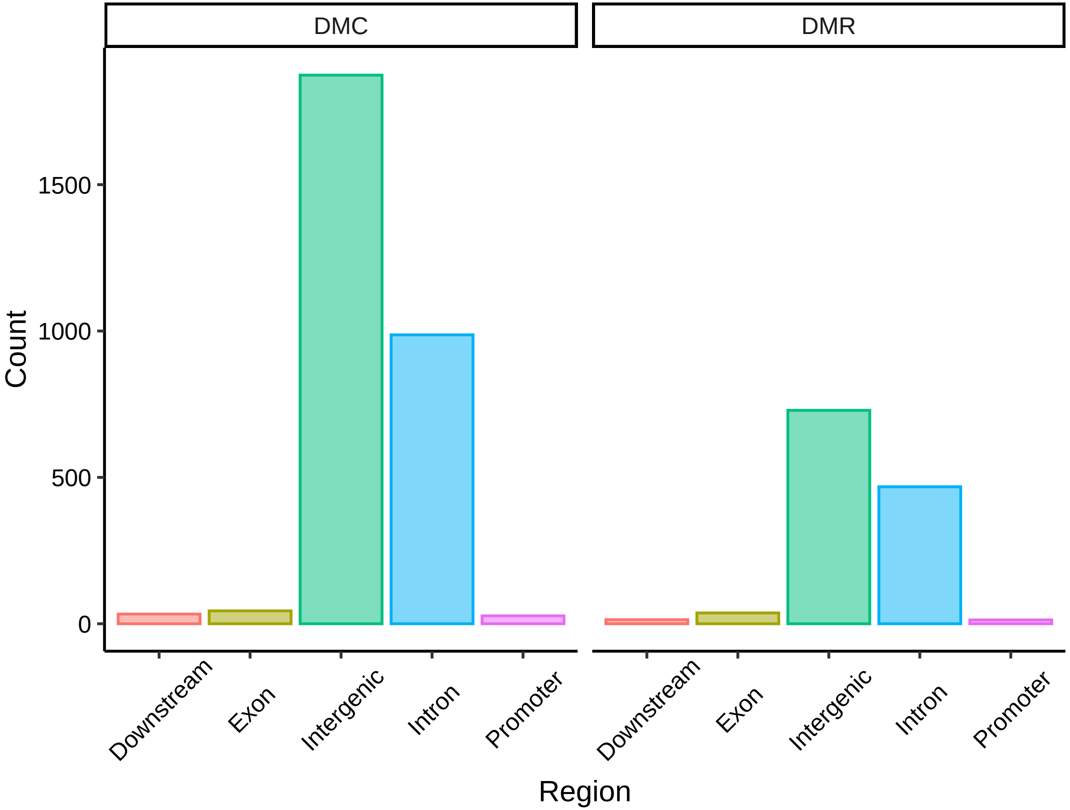
